# A Plasmid Network from the Gut Microbiome of Semi-isolated Human Groups Reveals Unique and Shared Metabolic and Virulence Traits

**DOI:** 10.1101/2021.08.30.458253

**Authors:** Liliane Costa Conteville, Ana Carolina Paulo Vicente

**Affiliations:** Laboratório de Genética Molecular de Microrganismos, Instituto Oswaldo Cruz, Fundação Oswaldo Cruz, Rio de Janeiro, RJ, Brasil

**Keywords:** Gut microbiota, Shotgun Metagenome, Traditional Groups, Antibiotic Resistance Genes, Metal Resistance Genes, Hunter-gatherers, Functional Analysis, T6SS

## Abstract

The plasmidome, the set of plasmids in gut microbiomes, has the potential to contribute to the microbiome assembly, as well as human health and physiology. Nevertheless, this niche remains poorly explored, likely due to the difficulties attributed to mining mobile elements from complex environments such as microbiomes. In general, most microbiome studies focus on urban-industrialized groups, but here, we studied semi-isolated groups, which represent a link between the ancestral and modern human groups. Based on metagenomic data, we characterized their plasmidome, including the set of accessory genes and functions from the gut microbiome of the Hadza, Matses, Tunapuco, and Yanomami. In silico analyzes revealed unique plasmid clusters and gene functions for each human group related to their diet and lifestyle. In addition, network analysis revealed a dozen plasmid clusters shared by these distinct groups but that are also circulating in other niches worldwide. Moreover, in these microbiomes, there are novel and unique plasmids associated with their resistome and virulome. A resistome encompassing six antibiotic classes and multiple metals, and a virulome with type VI secretion systems were identified. Functional analysis revealed pathways associated with urban-industrialized groups, such as lipopolysaccharide biosynthesis that was characterized in the Hadza plasmidome. These results demonstrate the richness of features in the semi-isolated human groups’ plasmidome and provide a snapshot of the forces that are acting in their gut microbiome. Their plasmidomes also represent an important source of information with biotechnological and/or pharmaceutical potential.

## Introduction

The human gut microbiome is a complex and dynamic ecosystem that performs several essential functions for human metabolism, nutrition, physiology, and immunity. Due to its high density and diversity of microorganisms, this environment is prone to constant genetic exchange between resident and transient bacteria^**1–4**^. Genetic transfers by plasmids in the gut have the potential to impact the ecology and evolution of the gut microbes. The acquisition of a plasmid and its cargo genes by a gut bacteria or archaea may assist the microorganism’s survival, fitness, and adaptation to the environment ^**4–7**^. Some traits provided by plasmids in the gut have been associated with antibiotic (AR) and metal resistance (MR)^**6,8–10**^, salt tolerance ^**11–12**^, and interbacterial competition^**13**^. Therefore, the human gut plasmidome has the potential to modulate the gut microbiome community and introduce new traits, such as antibiotic resistance and virulence to both resident and transient gut microbes^**4–14**^.

Several studies have assessed the bacterial profile present in human gut microbiomes^**15–19**^. Although these studies have improved our knowledge on the subject, the gut plasmidome is a niche in the gut microbiome that is still poorly investigated despite its potential to modulate the microbiome physiology and consequently, human health, in a one-step fashion. Many human aspects modulate the gut microbiome and plasmidome, including diet, lifestyle, therapy, consumption of pre- and probiotics, and diseases^**7,15–21**^. These aspects have been dramatically transformed during human history, which is evidenced by the current plethora of diets and lifestyles around the globe. Therefore, the gut microbiome composition and functionality are products of a long coevolutionary host-microbe relationship, and the human gut plasmidome also reflects this coevolution. As an example of the diet impacting mobile elements from the human gut, the seaweed in the Japanese daily diet was proposed to have transferred algal polysaccharide degradation genes to the Japanese microbiome^**20**^. The different diets of Fijians and Americans have been linked to the different glycoside hydrolases families encoded by mobile genes in the gut microbiome of these groups^**7**^. Moreover, the use of antimicrobials has been associated with the increase in the prevalence of mobile elements carrying functional resistance genes associated with synthetic antibiotics^**6–10,22–26**^.

Most of the microbiome studies have been focusing on urban-industrialized groups, and few have considered traditional human groups^**15–19,27**^. However, the study of gut plasmids from traditional groups has the potential to unravel genetic elements related to their unique lifestyle and environment. These groups retain valuable information about the microbiome before urbanization and industrialization impacted human diet and lifestyle, and also clues that can expand the field of prebiotics and probiotics for modern disorders prevention and treatment, as well as biomarkers that can form the basis for health and prognostic disease tests. So here, based on the gut microbiome from semi-isolated human groups, we investigated their plasmid content, including the set of accessory genes associated with these mobile elements that can be modulating human health and physiology.

## Results

We identified a total of 290 putative plasmids in the gut microbiome of the Yanomami/Brazil, Matses, Tunapuco, and Hadza. These plasmids were identified based on the nucleotide similarity with sequences of replication initiation protein (*rep*), mobilization protein (relaxase), mate-pair formation (MPF), and origin of transfer (*oriT*). The Yanomami/Brazil was the group with more plasmids identified (n=184), followed by the Hadza (n=52), the Matses (n=36), and the Tunapuco (n=18). The number of plasmids per metagenome varied from zero to 17 in the Yanomami/Brazil, Matses, and Hadza, but in the Tunapuco it varied from zero to seven plasmids. However, one Yanomami/Brazil metagenome stands out for harboring 90 plasmids (STable1).

The plasmids were classified as conjugative, mobilizable, and non-mobilizable based on three plasmid markers: relaxase, MPF, and *oriT*. Thus, among the 290 plasmids identified in this study, 8 (~ 2.7%) were classified as conjugative, for harboring both relaxase and MPF; 89 (~ 30.6%) as mobilizable, for harboring relaxase and/or *oriT* without MPF; and the majority (n= 193, ~ 66.5%) as non-mobilizable, for not presenting relaxase and *oriT*. Considering the plasmid length, most of them (79.6%) have less than 10Kb, with 20% having between 10 and 90 Kb, and the longest plasmid has 115 Kb. Information regarding each plasmid identified can be found in STable1.

Moreover, mobilization (MOB) typing was used to classify the relaxase and *oriT* sequences of the conjugative and mobilizable plasmids into six MOB types. MOBP was the most prevalent type, followed by MOBQ, MOBF **(Figure 1)**. Two other MOB families (MOBC and MOBV) were assigned to relaxases from the Hadza and Matses, respectively. MOBH was the only MOB family not identified. Besides it, 16 *oriT* sequences did not match any of the six known MOB families and were classified as unknown. The MPF was also classified into classes and the most prevalent was MPF_F_, followed by MPF_I_ and MPF_T_ **(Figure 1)**.

**Figure 1.**
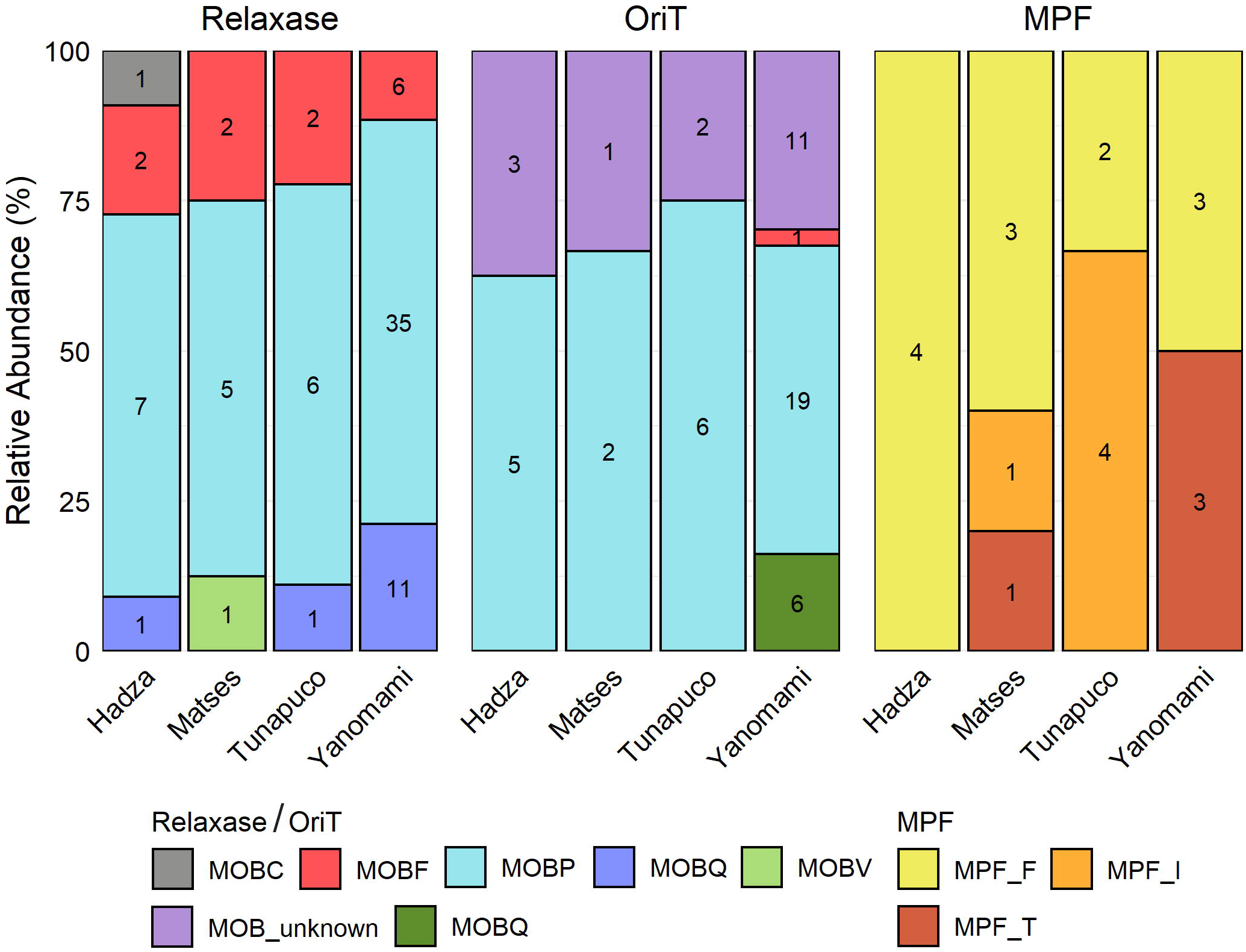
Stacked bar chart representing the relative abundance (y-axis) of the MOB families and MPF types of the relaxase, *oriT*, and MPF identified in the plasmidomes of each group. Raw counts are also showed.

We explored the protein’s functions encoded by the plasmid accessory genomes. The most abundant protein categories were Signaling/Cellular Processes (28-53%) and Genetic Information Processing (13-23%) (STable2). The majority of the proteins associated with Signaling/Cellular Processes are part of Secretion (IV and VI) and Prokaryotic Defense Systems. And the majority of proteins categorized as with Genetic Information Processing are related to Replication and Repair (STable2). Some relevant gene functions identified in the plasmidome differed by group. Functions related to lipopolysaccharide biosynthesis were exclusive to the Hadza, plant-pathogen interaction, immune system, and siderophore biosynthesis were exclusive to the Matses, flagellar assembly and drug and xenobiotics metabolism were exclusive to the Yanomami. Moreover, genes associated with Carbohydrate Metabolism and Other Amino Acids were only identified in the Matses and Yanomami. We also observed that the genes that generate energy metabolize different nutrients according to each group: Sulfur, Nitrogen and Methane in the Hadza, Matses and Yanomami, respectively.

Considering Prokaryotic Defense Systems, we performed focused analyses on identifying antimicrobial and metal resistance, and virulence, which in some cases, can directly impact the host health. AR and MR genes and virulence factors (VFs) were found in 27 plasmids identified in 19/72 metagenomes (STable1). The Hadza and the Yanomami/Brazil gut microbiomes are those with more AR, MR, or virulence genes. The AR genes identified are associated with six antibiotic classes: aminoglycosides, □-lactams, fluoroquinolones, fosfomycin, tetracyclines, and trimethoprim. The most abundant AR genes found were *aph* (aminoglycoside resistance), *tet* (tetracycline resistance), and *tem* (beta-lactamase). The MR genes are associated with copper, iron, mercury, and multiple metals resistance. Concerning the VFs, one plasmid presents a heat-stable enterotoxin (*astA*), but the other VFs are mainly yersiniabactin, which besides being a VF, is also associated with iron resistance. The gene *ybt*, which is a siderophore yersiniabactin, was identified in a cluster of eight *ybt* genes in one plasmid recovered from one metagenome from the Matses (STable1).

Based on the clustering classification, the plasmids were split into three groups: 1, 2, and 3. Group 1 comprises plasmids that were assigned to plasmid clusters of the MOB-suite program database, therefore known plasmids. While plasmids that exceeded the distance threshold with all references in the database were classified in Groups 2 and 3, and therefore novel plasmids could be part of these groups. The former comprises plasmids shared among the metagenomes and were clustered, and the latter comprises unique plasmids. The plasmids of Group 1 (n= 96) were grouped into 63 clusters, Group 2 (n= 65) into 22 clusters, while the remaining 129 were singletons (Group 3).

To infer any connection between the plasmids of the studied human groups, we generated a network based on the plasmid clusters **(Figure 2)**. The network revealed that 12/85 clusters (Group 1, n= 9; and Group 2, n =3) are shared among the human groups, mostly between the Hadza and the Yanomami. Interestingly, the Hadza plasmidome has counterparts with all other human groups. Moreover, to expand our scenario, we also performed BLAST analysis on the plasmids shared among the groups. In this way, we found that the semi-isolated groups’ plasmidome have high similarity with plasmids spread worldwide in bacteria from humans, animals, environments, and foods **(Table, STable3)**. Even Group 2 clusters, potentially comprising novel plasmids, presented similarities with known plasmids. However, the BLAST analysis of the plasmid cluster “R” returned no matches, indicating the presence of novel plasmids in this cluster.

**Figure 2.**
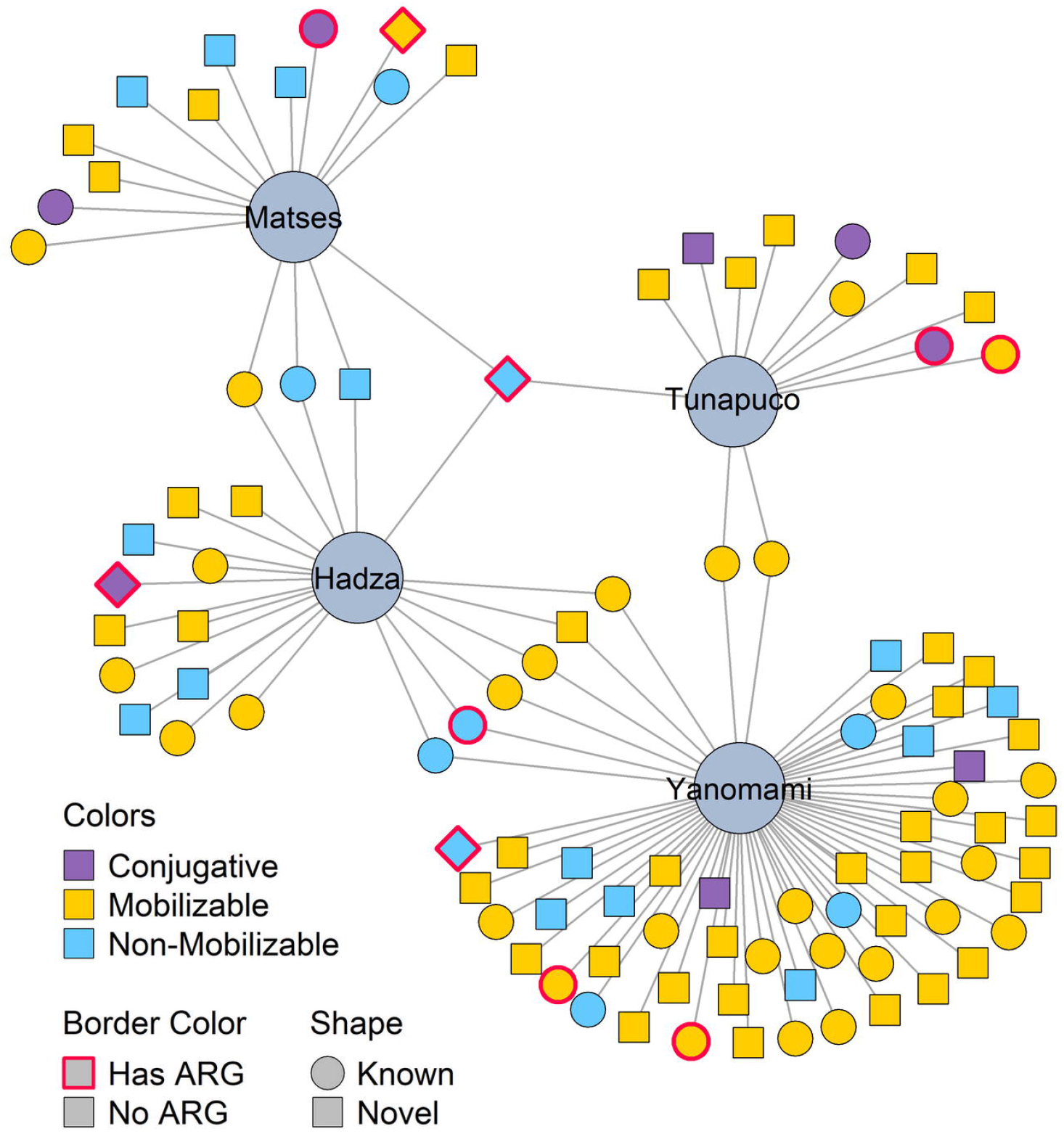
Network based on the plasmid clusters (smaller-sized nodes) harbored and shared between the human groups (bigger-sized nodes). Non-mobilizable plasmids that were identified in only one metagenome were removed from this analysis.

Surprisingly, the plasmids from the clusters “A”, “E”, and “C” **(Figure 2)**, previously characterized as non-mobilizable **(STable1)**, were the clusters identified in a higher number of metagenomes. The cluster “A” was the most prevalent in the metagenomes analyzed, present in the Hadza (n=3), Matses (n=3), and Tunapuco (n=1). The same *rep* and *TolA* sequences characterized the plasmids from cluster “A”, including the Hadza plasmids (n=3), which also harbor a *dfrf* (dihydrofolate reductase) trimethoprim resistance gene. Besides that, this cluster was inferred by the MOB-suite as potentially hosted by bacteria of the genus Campylobacter. The clusters “E” and “C” harbor known plasmids, with no relevant functions so far assigned. The mobilizable cluster “F” shared between the Hadza and Yanomami carries beta-lactamase resistance genes, besides some Hadza’s plasmids also carry *aph* (aminoglycoside resistance). Conjugative plasmids carrying AR genes were identified in cluster “M” (*fosA* gene), cluster “O” (*aph* and *tet* genes), and cluster “P” (*ctx* gene) **(Figure 2)**.

Besides the plasmids shared among the human groups, we also observed plasmids with the potential to be human group-specific as some from Group 2 clusters (n = 8) identified only in metagenomes from Hadza (n=2) and Yanomami (n=6). The Yanomami-specific clusters were classified as mobilizable (n=4), but the others and Hadza’s plasmids were non-mobilizable. These “human group-specific” plasmid clusters carry genes associated with Metabolism, mostly related to Glycolysis/Gluconeogenesis, and also genes that encode Transport Proteins and Transposases.

## Discussion

Diet and lifestyle have changed dramatically throughout human evolution, first with the introduction of farming (~ 10,000□years ago), and more recently, with the introduction of industrially processed foods, hygienic conditions, antimicrobials, pollution, and other factors^**27,28**^. These transformations in human lifestyle have been widely associated with the selection and modulation of the gut microbes and with a substantial loss of gut bacterial diversity ^**15–19,27**^. However, this microbial profile restructuration does not only involve the microbial taxa but, responding to selective pressures, also the mobilome^**4**^. Moreover, modern diseases, such as diabetes, obesity, and allergies, partially driven by the lifestyle, have been associated with the loss and variation of bacterial taxa and mobile elements diversity in the gut microbiome^**28,29**^.

In the present study, we analyzed the gut plasmidome and its accessory genes of four semi-isolated human groups: the Yanomami from the Brazilian Amazon Forest^**19**^, the Matses from the Peruvian Amazon Forest, the Tunapuco from the Peruvian Andean highlands^**16**^, and the Hadza from northwestern Tanzania^**17**^, whose gut microbiomes represent a link between the ancestral and modern human groups. The bacterial composition and functionality of these four human groups have been previously analyzed, and although they have a similar contrasting composition compared to urban-industrialized groups, each of them has particular characteristics related to their different ecological niches, genetics, and diets^**15–17,19**^. The plasmidome analysis showed a similar pattern, in which specific plasmid clusters and gene functions are unique for each human group, yet they also share significant parts of the plasmidome. Unexpectedly, a dozen of plasmid clusters are shared among these semi-isolated human groups living in distinct geographical niches. In addition, plasmids from the shared clusters have also been detected worldwide in humans, animals, environments, and foods. This reveals that distinct semi-isolated human groups and their environments have been impacted by the same modern practices eventually carried out by worldwide urban-industrialized individuals.

In general, plasmids can be classified by their relaxase and *oriT*, which comprise six MOB families, and MPF, which comprises four families^**23,30**^. Based on a database of plasmid markers, the plasmidomes of our study were mostly characterized by the families MOBP, MOBQ, and MOBF. The MOBP, a cluster of actively evolving relaxases, is the most prevalent family in plasmids databases, characterizing a considerable proportion of short plasmids^**30,31**^. Indeed, most of the plasmids that we identified are short plasmids (<10Kb). Moreover, the MOBC and MOBV families, low prevalent families in plasmid databases, were identified in two novel plasmids of the Hadza and Matses. Revealing the yet underexplored scenario that modulates the gut plasmidome and microbiome diversity^**30,31**^. Concerning the quantitative aspect, the Yanomami presented the largest plasmidome and the Tunapuco the smallest, likely reflecting their remarkably different diet, lifestyle, and environment. Although both groups inhabit the Amazon Region, the Yanomami’s subsistence is based on gathering and hunting wild plants and animals in the Amazon Forest, the world’s largest and most biodiverse tropical rainforest^**19,32,33**^. In contrast, the Tunapuco’s subsistence is based on local agricultural produce and homegrown small animals located in the Peruvian Andean highlands, at an elevation between 2,500 and 3,100 m above sea level^**16,32,33**^.

The gut microbes usually process undigested carbohydrates, proteins, and fat, producing additional energy from the diet and also metabolites that can impact human physiology^**34**^. These metabolic functions, encoded by chromosomes and plasmids of the gut microbes, are selected/modulated according to the components of the human’s host diet^**7,20,34,35**^. In these groups, the food seasonality and their eventual nomadic behavior are selection pressures that can drive a variety of changes in the gut throughout the year, particularly in the plasmidome. So, to rapidly adapt to the new/seasonal dietary components, specific enzymes must be selected and spread among the gut microbes^**34,35**^. This may explain why only the groups that inhabit the Amazonian Forest (Yanomami and Matses) have plasmid accessory genes associated with Carbohydrate and Other Amino Acids Metabolism. These groups live in a hotspot of diversity, therefore they are always coming into contact with various dietary components, which must be metabolized^**32–35**^. Moreover, their niches and consequently, the distinct selective pressures, contribute to the selection of plasmids’ content that participates in the metabolism of molecules from their diet. The functions identified as group-exclusive follow this context. For example, only in the Hadza plasmidome was observed the lipopolysaccharide (LPS) biosynthesis pathway. This pathway is known for inducing strong inflammatory responses, but contrastingly, it also has anti-inflammatory actions when produced by certain gut bacteria, such as Bacteroidetes. Indeed, the Hadza harbor this phylum in their gut microbiomes^**15,17**^. This LPS has been proposed to be used in the prevention or treatment of chronic inflammatory diseases that are induced by pro-inflammatory LPS^**36**^. Therefore, the gut accessory genome, represented by the plasmidome, constitutes an underexplored plethora of genes and functions and an important source of information with potential biotechnological or pharmaceutical value^**37,38**^.

The prevalent gene functions that were identified in the plasmidome of all groups, such as Prokaryotic Defense Systems, Secretion Systems, and Replication and Repair, are associated with core plasmid genes^**39,40**^. However, the type VI Secretion Systems (T6SS) is not plasmid ubiquitous, moreover, it is associated with interbacterial competition, bacteria-host interactions, and virulence^**41–43**^. Therefore, if the T6SS is harbored by a commensal microbe, it can prevent the microbiome disassembly by an invading bacteria. In contrast, if T6SS is carried by an invading bacteria, the community assembly can be disestablished, resulting in dysbiosis^**41–43**^.

Interestingly, we observed a resistome in the plasmidomes of these human groups that have no or limited exposure to synthetic antibiotics, as well as reported in other isolated or semi-isolated groups/locations^**7,24,44**^. Most of these AR genes are associated with β-lactam and tetracycline resistance, which are commonly seen in soil and water environments^**45**^. Therefore, the AR genes in these semi-isolated groups may have originated from the environment where antibiotic-producing microbes naturally occur, and also may have reached these groups via contacts with antibiotic-exposed populations^**22,24,46,47**^. In fact, Hadza and Yanomami plasmidome harbor accessory genes with the function of biosynthesis of Penicillin and Cephalosporin. In the same way, metal resistance genes present in the plasmidome can reflect bacterial adaptations to environmental conditions. Indeed, the Yanomami niche is likely to be contaminated with cadmium due to the continuous discharge of batteries by them over decades^**19**^, however, the natural or artificial presence of metal in the other group’s environment is unknown. Moreover, MR genes can impact the community assembly of the gut microbiomes and consequently affect host health^**48**^. Concerning the virulome, there is no prevalence of bacterial virulence-associated factors in the plasmidome, with the expectation of T6SS. In fact, these factors are not prevalent within environmental bacteria but occur more often in pathogens in clinical settings^**49**^.

The plasmidomes recovered and characterized here represent an achievement in the study of the gut microbiome of semi-isolated groups, providing a snapshot of the forces that modulate the gut microbiome given the unique as well as global scenarios to which these human groups are exposed. Although we have focused only on plasmids from the human gut, other mobile elements, such as bacteriophages, must also be relevant in this context. Beyond that, we are aware that our study has biases and limitations, for example, the plasmid identification that is based on the current plasmid markers in the databases. The semi-isolated groups may harbor plasmid markers that have not yet been characterized and this may have resulted in the general lack of relaxase, MPF, and oriT sequences identified. As a result, most plasmids were classified as novel and non-mobilizable. Indeed, new MOB families have often been reported^**30,50–54**^. Besides, the complexity of metagenomic assembly tends to produce shorter contigs and may have hampered the identification of the whole genetic content of the plasmids^**55**^. In addition, the largest number of genes found in the plasmids have no known functionality. Indeed, the gut plasmidome is a recognized source of novel genes and gene products, so the presence of unknown or novel elements is expected^**38**^. However, these results highlight the knowledge gap in this field and in databases, which is often attributed to the difficulties in studying plasmids from complex environments, as microbiomes^**40,56**^.

## Materials and Methods

For this study, we analyzed shotgun metagenomic data from previously published gut microbiome studies of traditional human groups: semi-isolated hunter-gatherer communities from South America and Africa (Yanomami from Brazilian Amazon^**19**^, n = 15; Matses from Peruvian Amazon^**16**^, n = 24; and Hadza from Tanzania^**17**^, n = 27), and a rural agricultural community from the Andean highlands in Peru (Tunapuco^**16**^, n = 12).

Each metagenome was independently subjected to de novo assembly through metaSPAdes v.3.13^**57**^. For plasmid identification, MOB-recon and MOB-typer modules from the MOB-suite program were used^**58**^. These modules identified putative plasmids into the contigs of each metagenome and searched for replication gene (*rep*), mobilization protein (relaxase), matepair formation (MPF), and the origin of transfer (*oriT*). Based on the presence/absence of these plasmid markers, the plasmids’ mobility was predicted^**58**^. To avoid false positives we removed from the analysis the plasmids with no markers identified by the MOB-suite. For characterization, pairwise genomic distances between each plasmid identified and each reference plasmid cluster was calculated. Thus, each plasmid identified was assigned to their closest reference plasmid cluster or as a novel plasmid depending on the predicted genomic distance^**58**^. Following the MOB-suite analysis, the function “dist” of the Mash program^**59**^ was used to calculate the pairwise genomic distances between all novel plasmids identified in the groups. If the distance between the novel plasmids was lower than 0.05, they were assigned to the same novel cluster. This distance metric allows the clustering of plasmids with considerable differences in length, which is useful in this case, since plasmids can undergo diverse changes in their sequence content^**58**^. Based on the clustering classification, the plasmids were split into three groups. Group 1 comprises plasmids that were assigned to plasmid clusters from the reference database. Group 2 comprises plasmids assigned as novel and clustered into novel clusters. Group 3 comprises unique plasmids, with no similarity to the plasmids in the database or to other plasmids found in the metagenomes. Moreover, only plasmid clusters identified in at least three metagenomes from one same human group were considered with the potential to be group-specific.

We also used FragGeneScan v.1.31^**60**^ to predict Open Reading Frames (ORFs) in the contigs assigned as plasmids. Thereafter, Abricate (https://github.com/tseemann/abricate) was used to screen the ORFs against the MegaRes v.2^**61**^ and VFDB^**62**^ databases. Abricate results were filtered for a minimum DNA identity of 75% and minimum coverage of 50%^**63**^. Functional annotation and assignment of KEGG functional categories to the ORFs were performed using the KOfamKOALA tool^64^, considering a threshold of 0.01.

Network analysis of the plasmids identified in the four groups was performed based on the plasmid clusters characterized by MOB-suite. The network was constructed with the igraph R package (http://igraph.org/r/). In this network, the smaller-sized nodes represent the plasmids, which are linked to the bigger-sized nodes that represent the groups that harbor these plasmids. We removed from this analysis non-mobilizable plasmids that were identified in only one metagenome.

In order to investigate which other hosts and niches these plasmids have been identified in, we performed a search analysis using the online BLAST tool^**65**^ on each plasmid. We removed results with similarity and coverage less than 70% and e-value less than 1e-5. To recover metadata information from the accession numbers of the plasmids that matched our putative plasmids, we used a custom python script based on the Entrez module of the Biopython package.

## Data availability statement (DAS)

The metagenomic data of the gut microbiomes used in this study were downloaded from the National Center for Biotechnology Information Sequence Read Archive (BioProject: PRJNA268964 for Reference 16, PRJNA278393 for Reference 17, and PRJNA527208 for Reference 19).

## Table Legend

Table. Mobility prediction and Antibiotic Resistance Genes of the plasmid clusters A-T from the network. The hosts and niches shown refer to the plasmids that had high similarity to this cluster by blast analysis. Groups: H - Hadza; M - Matses, T - Tunapuco, Y - Yanomami

